# Minimizing carry-over PCR contamination in expanded CAG/CTG repeat instability applications

**DOI:** 10.1101/187872

**Authors:** Loréne Aeschbach, Vincent Dion

## Abstract

Expanded CAG/CTG repeats underlie the aetiology of 14 neurological and neuromuscular disorders. The size of the repeat tract determines in large part the severity of these disorders with longer tracts causing more severe phenotypes. Expanded CAG/CTG repeats are also unstable in somatic tissues, which is thought to modify disease progression. Routine molecular biology applications involving these repeats, including quantifying their instability, are plagued by low PCR yields. This leads to the need for setting up more PCRs of the same locus, thereby increasing the risk of carry-over contamination. Here we aimed to reduce this risk by pre-treating the samples with a Uracil N-Glycosylase (Ung) and using dUTP instead of dTTP in PCRs. We successfully applied this method to the PCR amplification of expanded CAG/CTG repeats, their sequencing, and their molecular cloning. In addition, we optimized the gold-standard method for measuring repeat instability, small-pool PCR, such that it can be used together with Ung and dUTP-containing PCRs, without compromising data quality. We expect that the protocols herein to be applicable for molecular diagnostics of expanded repeat disorders and to manipulate other tandem repeats.

## Introduction

Tandem trinucleotide repeats (TNRs) are common in the human genome and are polymorphic in the population^1^. Their size at any one locus is usually below about 35 triplets. When these sequences expand beyond this threshold, however, they become highly unstable with a bias towards expansions in both somatic and germ cells^2-5^. In addition, TNR expansions cause a wide variety of human neurological, neuromuscular, and neurodegenerative disorders^4^. Importantly, longer repeat tracts tend to provoke more severe symptoms^6^. Therefore, expansions from one generation to the next leads to an earlier age of onset in the progeny, providing an elegant molecular explanation for the phenomenon of anticipation. Meanwhile, ongoing expansions in somatic tissues may accelerate pathogenesis^4,7-9^.

The mechanisms of repeat instability remain tedious to assay in part because of the low yields during PCR amplification of expanded TNRs^5^. This is especially true for CAG/CTG repeats (henceforth referred to as CAG). Thus, several PCRs are often setup in parallel to obtain sufficient amounts of DNA for downstream applications. This opens the door to carry-over contamination in which the amplicon produced in a previous reaction contaminates the subsequent ones, skewing the results or rendering them useless. Typical methods to palliate this risk include having a dedicated ‘pre-PCR’ room, pipettes, and solutions. A hood with a UV light source can also be used. These precautions reduce, but do not eliminate, the risk of contamination. This is particularly important in the context of small-pool PCR (SP-PCR), the gold standard assay to measure repeat instability^10,11^. This technique consists of performing many PCRs in parallel, each using only a few genomes as template to bypass the propensity of amplifying the shorter alleles in the population. The products are then separated on agarose gels and transferred onto a membrane for Southern blotting. This assay is exquisitely sensitive and thus many no-DNA controls must be included to ensure that there is no carry-over contamination. If the controls show amplification products, the experiment must be repeated with fresh solutions, wasting time and resources.

Carry-over contamination is a problem for molecular diagnostics^12^, ancient DNA amplification^13^, and for any application that requires amplifying the same few genomic loci^14^. A common solution is to pre-treat the samples with a bacterial uracil *N-*glycosylase, Ung, that degrades any contaminating DNA containing uracil and to replace dTTP with dUTP in all PCRs^15^. It was unclear whether this method is suitable for molecular applications involving expanded CAG repeats given the difficulties associated with their PCR amplification. Here we successfully adapted protocols and show the applicability of this method in four molecular biology techniques commonly used when working with expanded CAG repeats: PCR amplification, sequencing, molecular cloning, and SP-PCR.

## Results and Discussion

### dTTP substituted with dUTP allows PCR amplification of expanded CAG repeats

We first wanted to determine whether expanded CAG repeats of varying lengths could be amplified using dUTP instead of dTTP in the PCR and that Ung did not interfere with the amplification. We therefore pre-treated samples with Ung and set up PCRs containing either dUTP and dTTP (Fig. 1A). We used genomic DNA isolated from HEK293-derived cells harbouring a single copy transgene with repeat sizes ranging from 15 to 101 (GFP(CAG)x) ^16^. We found that the Ung-treated and dUTP-containing reactions consistently had a slightly lower yield for repeat sizes ranging from 15 to 101 CAG repeats compared to dTTP-containing reactions (Fig. 1B). This did not affect the apparent size of the repeat tract after agarose gel electrophoresis. Importantly, uracil-containing amplicons were efficiently degraded by Ung prior to PCR (Fig. 1CD). We conclude that pre-treatment with Ung and the use of dUTP in the PCR allows for an efficient amplification of expanded CAG repeats and prevents carry-over contamination.

**Figure 1:**
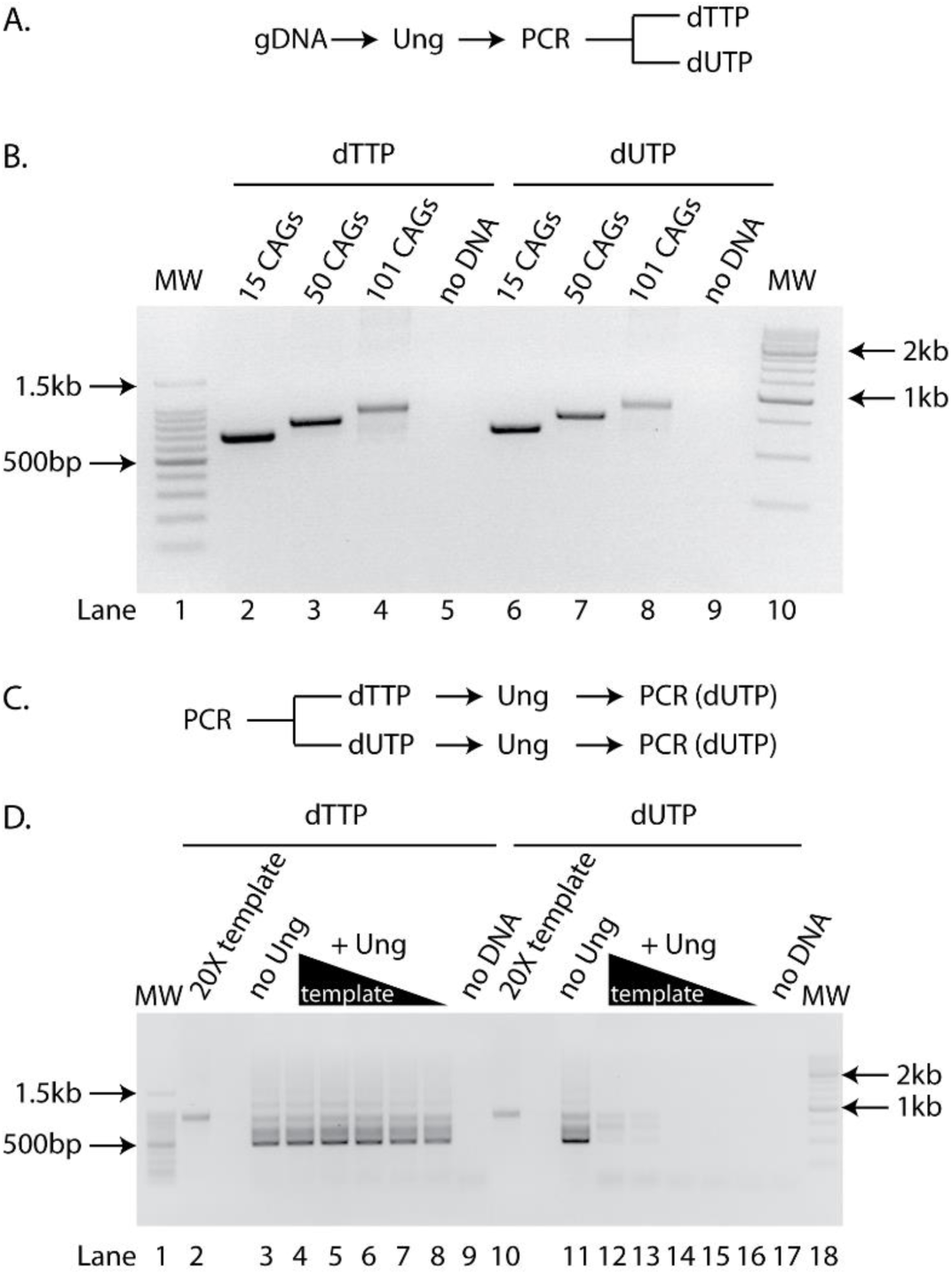
dTTP substituted with dUTP allows PCR amplification of expanded CAG repeats. A) Experimental scheme used to generate the PCR amplicons found in B. B) Agarose gel of PCR amplicons after pre-treatment with Ung and containing either dTTP (lanes2-5) or dUTP(lanes6-9)from genomic DNA isolated from GFP(CAG)_15_ (lanes 2, 6), GFP(CAG)_50_ (lanes 3, 7), GFP(CAG)_101_ (lanes 4, 8), or without DNA added (lanes 5, 9). MW = molecular weight marker (lanes 1, 10). C) Experimental scheme used for panel D. D) Agarose gel of PCR amplicons obtained after pre-treatment with Ung of a template containing either dTTP (lanes 2-9) or dUTP (lanes 10-17). The template was generated from amplifying GFP(CAG)_101_ genomic DNA. The PCRs with no Ung treatment (lanes 3,11) contained 1X concentration of the template amplicon. Ten-fold dilution series (lanes 4-8 et 12-16) of the template (20X loaded directly without PCR in (lanes 2, 10)) and no DNA controls are also shown (lanes 9 and 17). MW = molecular weight marker(lanes1,18).

### dUTP-containing amplicons can be sequenced with ease

We next determined whether the amplicons containing uracil could be sequenced as efficiently as those containing thymine. To do so, we set up 6 different sequencing reactions from two PCR products amplified with either dUTP or dTTP from genomic DNA obtained from GFP(CAG)_101_ cells. We found no difference in the apparent size determination between the two nucleotides used (Table 1, P=1.0 between uracil and thymidine-containing amplicons using a Wilcoxon U-test). These observations show that dUTP can be substituted for dTTP when sequencing of expanded CAG repeats.

**Table 1:**
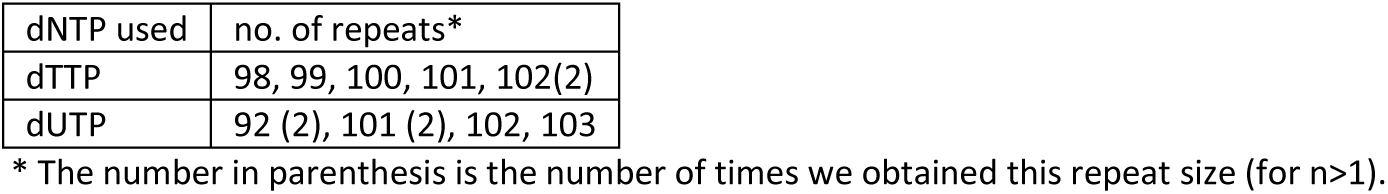
dUTP and dTTP yield the similar repeat sizes upon Sanger sequencing

| dNTP used | no. of repeats* |
| --- | --- |
| dTTP | 98, 99, 100, 101, 102(2) |
| dUTP | 92 (2), 101 (2), 102, 103 |

\* The number in parenthesis is the number of times we obtained this repeat size (for n>1).

### Keeping residual activity of Ung in check

We noticed that the Ung-treated and uracil-containing amplicons degraded completely at room temperature over a period of 3 days (Fig. 2). We saw some degradation even at 4°C, but none when the samples were stored at -20°C. This is consistent with the observation that Ung retains enzymatic activity even after many PCR cycles^17^. Indeed, the Ung that we used is active at temperatures below 55°C even after repeated denaturation-renaturation cycles, according to the manufacturer. Given that some applications, such as SP-PCR and molecular cloning, require small amounts of DNA, degradation may have a large impact. We circumvented this problem using two different strategies. The first was to deactivate the Ung by digesting with Proteinase K immediately after the reaction. The second approach was to use a heat-labile (hl) Ung^18^ (Fig. 2). This enzymes works well at 20°C, but an incubation at 95°C for 7 minutes inactivates it.

**Figure 2:**
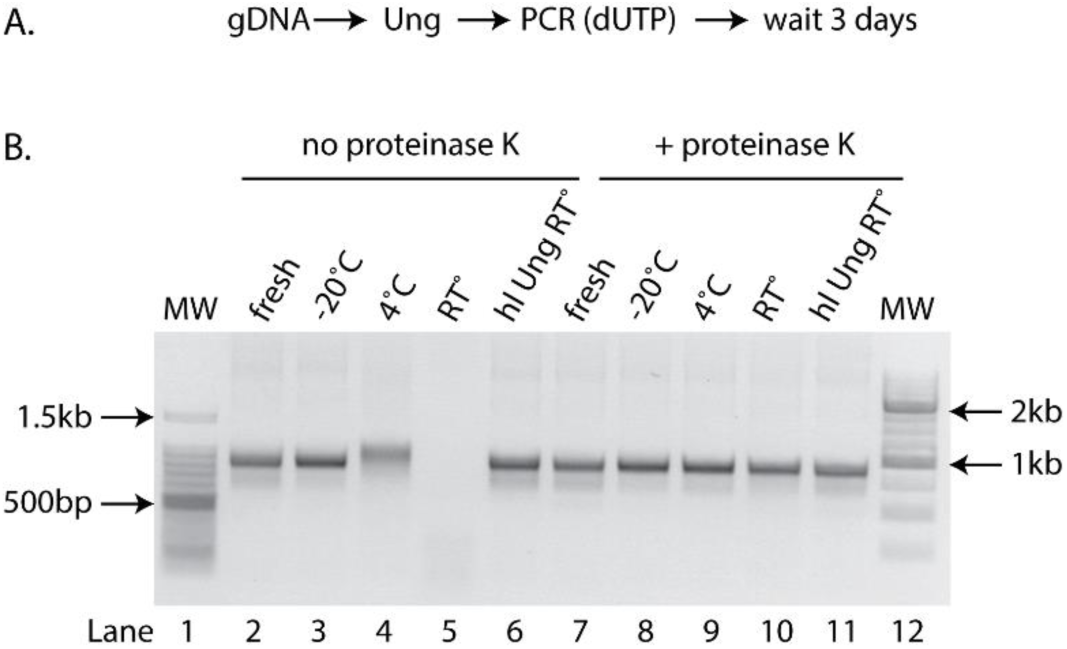
Using heat-labile Ung and/or Proteinase K prevents the degradation of uracil-containing amplicons. A) Experimental scheme used in panel B. B) Agarose gel of amplicons from genomic DNA of GFP(CAG)1_0_1 containing uracil and treated post-PCR with (lanes 7-11) or without (lanes 2-6) proteinase K three days after the PCR was performed (lanes 3-6 and 8-11) or immediately after the PCR (fresh, lanes 2, 7). The samples were stored at -20°C (lanes 3, 8), 4°C (lanes 4, 9), room temperature (RT°, lanes 5, 10), or were amplified using a a heat labile (hl) Ung used instead of the conventional Ung and stored at room temperature (lanes 6,11). MW = molecular weight marker (lanes 1 and 12).

### Cloning dUTP-containing expanded repeats

We next sought to determine whether it is possible to clone uracil-containing amplicons reliably. This was previously done using *ung^-^ dut^-^ E. coli* such that the plasmids isolated from these strains contained up to 20% of uracil^13^. Wild-type strains, by contrast, have less than one uracil residue per 10^6^ nucleotides^19^. Here we wanted to know whether we could clone uracil-containing amplicons using a *ung*^-^ strain and convert uracil to thymine residues in the process. Thus, we knocked out *ung* in DH5α cells. This strain background is used for a variety of molecular cloning purposes. We used the well-established lambda Red recombination-based approach^20^,^21^ to replace the endogenous *ung* with the coding sequence of a gene that provides resistance to spectinomycin, generating a *Δung* allele, *ung*^-^9. Thus, the more commonly used antibiotics, including ampicillin, chloramphenicol, and kanamycin,, remain available.

We tested whether cloning with uracil-containing amplicons was possible by amplifying a non-repetitive sequence, the SUMO1 cDNA, and then cloning it using the Zero Blunt TOPO PCR kit. Significantly, we were unsuccessful in cloning uracil-containing amplicons when we used the standard Ung during the pre-PCR treatment (data not shown). This was presumably because of degradation of the amplicons prior to transformation. We overcame this problem by using the hlUng before and proteinase K treatment after the PCR. Under these conditions, we found that cloning was efficient with thymine-containing amplicons in both *ung^+^* and *ung*^-^ strains (Table 2). By contrast, cloning uracil-containing amplicons with the *ung^+^* strain yielded no colony, presumably because the uracil residues are digested and the plasmid degraded once inside the cell. In *ung*^-^ cells, however, the cloning efficiency was similar whether uracil or thymine was present in the amplicon (Two-tailed Student’s t-test p=0.12). We next performed colony PCR and/or Sanger sequencing of the plasmids from 8 independent clones from each of the conditions that yielded colonies. In every case, we could confirm the presence of the SUMO1 cDNA insert (not shown). Taken together, our results demonstrate that it is possible to clone uracil-containing amplicons in *ung*^-^ bacteria.

**Table 2:** Number of colonies obtained when cloning thymine or uracil-containing amplicons of the SUMO1 cDNA

| Nucleotide used in PCR | DH5α* | DH5α *Δung* * |
| --- | --- | --- |
| dTTP | 60 ± 49 | 200 ± 34 |
| dUTP | 0 ± 0 | 142 ± 26 |
| No insert† | 0 ± 0 | 0 ± 0 |

†: The cloning reaction was performed in the absence of any insert to test for self-circularization of the plasmid backbone.

To determine whether our *ung*^-^ strain could convert uracil to thymine residues, we isolated plasmids from both *ung^+^* and *ung*^-^ strains and digested them with Ung *in vitro.* We found no evidence that plasmids isolated from our *ung*^-^strain contained any uracil (Fig. 3), suggesting that these residues are indeed converted to thymine *in vivo.*

**Figure 3.**
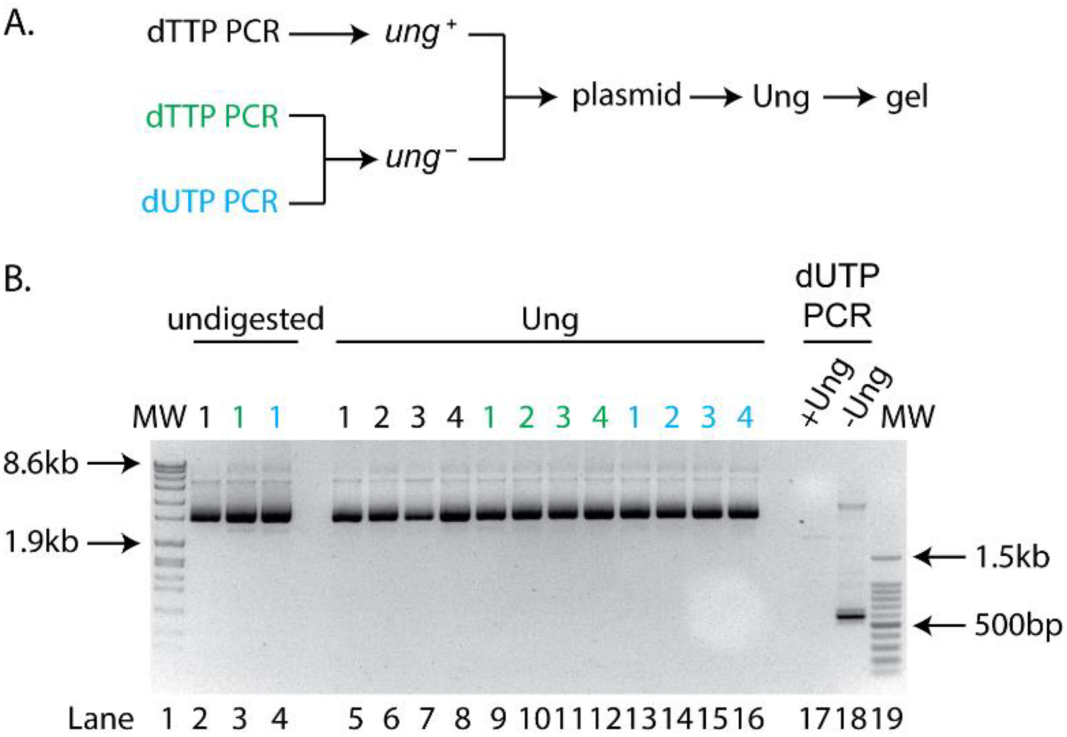
Plasmids cloned into *ung*^-^ contain undetectable levels of uracil. A) Experimental scheme used. B) Gel of plasmids isolated from *ung*^+^ cells transformed with a thymine containing insert (black – lanes 2, 5-8) or *ung*^-^ cells transformed with dTTP (green – 3, 9-12) or dUTP (blue – 4, 13-16). The samples remained untreated (lanes 2-4) or were incubated with 0.5U of Ung (lanes 5-16). Controls for Ung digestion consist of SUMO1 cDNA amplicons containing uracil digested with Ung (lane 17) or left untreated (lane 18). MW = molecular weight markers (lanes 1,19). The numbers above the lanes refer to the name of the clones tested.

We considered two important parameters when cloning expanded repeats. First, the efficiency of the cloning reaction itself and, second, the instability incurred during the cloning procedure. To assess the former, we used the same approach as for cloning the SUMO1 cDNA and calculated the number of colonies obtained when cloning a PCR product amplified from a plasmid sporting an unstable repeat tract of up to 89 CAGs and containing uracil or thymidine into *ung^+^* and *ung*^-^ strains (Table 3). We found similar numbers as for the SUMO1 cloning: uracil-containing amplicons only survived in the strain lacking *ung.* These results suggest that cloning uracil‐ and expanded CAG-containing amplicons can be achieved at high efficiencies in a *ung*”strain.

**Table 3:** Number of colonies obtained when cloning thymine or uracil-containing amplicons with 89 CAGs

| Nucleotide used in PCR | DH5α* | DH5α *Δung* * |
| --- | --- | --- |
| dTTP | 193 ± 117 | 88 ± 59 |
| dUTP | 0 ± 0 | 54 ± 45 |
| No insert† | 0 ± 0 | 0 ± 0 |

\*: number of colonies derived from 3 independent transformations of 3 independent PCR amplicons. The competence was similar with 10<sup>8</sup> colony/μg of pUC19 for both strains. The same amount of DNA was used for the cloning and transformation reactions (see methods).
†: The cloning reaction was performed in the absence of any insert to test for self-circularization of the plasmid backbone. The no insert experiments in the *Δung* were done twice.

To assess whether uracil-containing amplicons retained the expanded repeats throughout the cloning procedure, we isolated plasmid DNA from 10 different colonies from all three conditions yielding colonies and counted the number of CAGs blindly after Sanger sequencing. As expected, the plasmids isolated contained between 5 and 64 CAGs from the plasmids isolated, all contained the repeat tract. Interestingly, we found that the size distribution was slightly longer when we transformed in a *ung*^-^ strain uracil-containing inserts compared to thymine-containing ones (Fig. 4 - P= 0.03 Wilcoxon Utest), implying that *ung*–strains and uracil-containing amplicons are conducive to cloning expanded repeat tracts.

**Figure 4:**
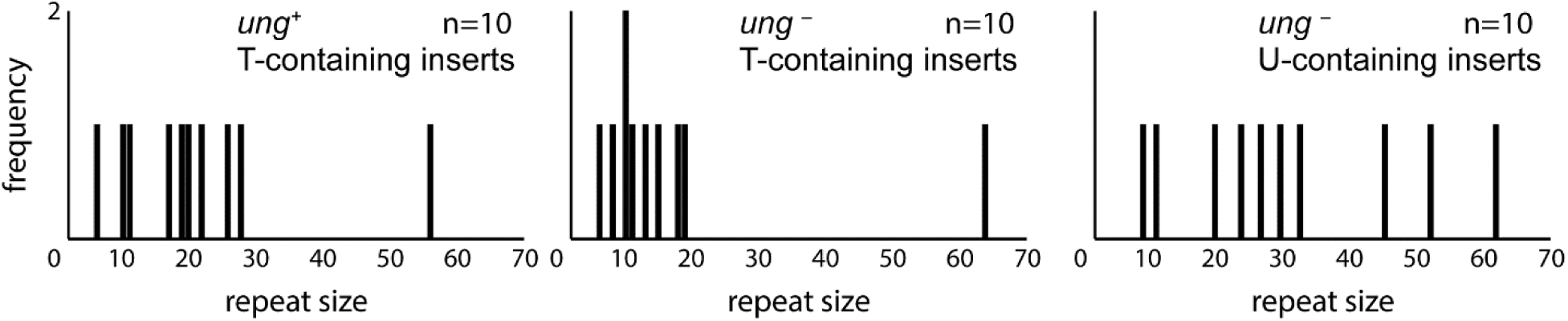
Repeat sizes recovered after molecular cloning of amplicons containing uracil or thymidine using DH5α and DH5α *Δung* strains.

### Small-pool PCR using Ung pre-treatment and dUTP

We next asked whether the gold standard assay to measure repeat instability frequencies, SP-PCR, was amenable to Ung treatment and dUTP-containing PCR. We expected this assay to be a challenge because of the small amounts of DNA used as templates and of the residual activity of Ung may lead to degradation of the products before we can visualize them. Indeed, we found that using the standard Ung led to blots so smeary that the results were unusable, which implied a considerable level of degradation. The treatment with proteinase K decreased the smearing somewhat (Fig. 5A). We observed sharp bands on the membranes only once we combined proteinase K and hlUng (Fig. 5B). These SP-PCR blots were done using a probe amplified after a Ung pre-treatment and containing dUTP, showing that the presence of uracil does not impair nick translation. We conclude that with these minor tweaks to the standard protocols, SP-PCR can be done with hlUng and dUTP.

**Figure 5:**
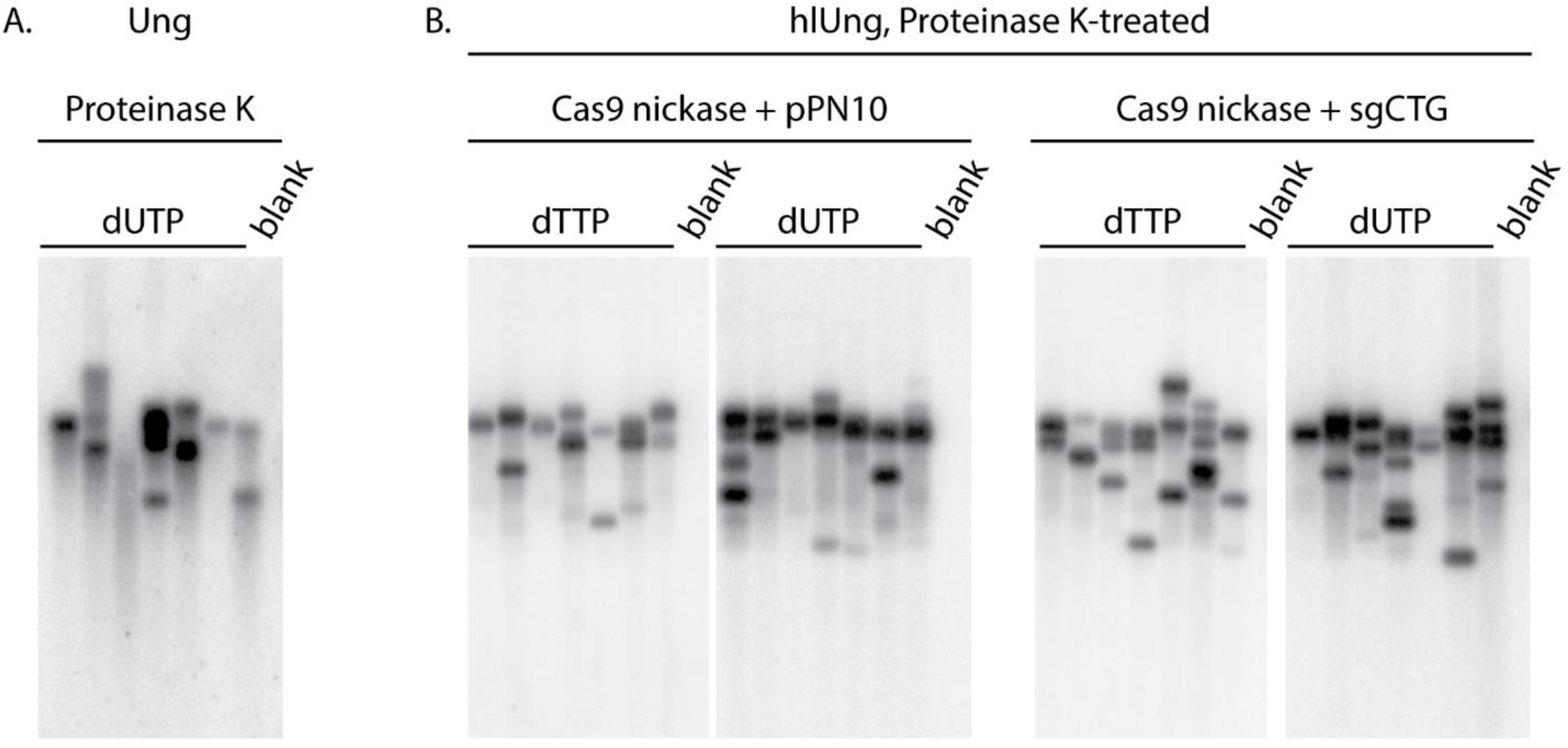
Small-pool PCR of expanded repeats using Ung and dUTP. A) SP-PCR blot from reactions pretreated with the standard Ung, amplified with dUTP, and stopped using Proteinase K. The smearing below the bands is evident. Each lane contains the product of a PCR with 50 pg of template DNA derived from GFP(CAG)_101_ treated with sgCTG and the Cas9 nickase. B) SP-PCR blots of amplicons generated using dTTP or dUTP, pretreated with hl Ung, and stopped after the PCR with Proteinase K. Each lane contains the product of a PCR that used 100 to 250 pg of genomic DNA from GFP(CAG)_101_ treated with the Cas9 nickase and either the sgCTG or an empty guide RNA vector (pPN10). In all cases, the probe used was generated with dUTP.

We confirmed that neither the hlUng treatment nor the use of dUTP affect the accuracy of SP-PCR by performing the assay on GFP(CAG)_101_ cells that were treated with the Cas9 nickase and a single-guide RNA targeting the CAG tract^16^. We found previously that this treatment led to an increase in the number of contractions found in the sample. Indeed, we observed here the increase in the number of contractions in both the SP-PCR made with dTTP and dUTP (Fig. 5B). These results demonstrate that the measures used to prevent carry-over contamination do not affect the result interpretation.

Our experiments show that a pre-treatment with Ung and the use of dUTP in PCRs are compatible with the most widely used molecular biology techniques involving expanded CAG repeats. In particular, we can PCR-amplify, sequence, and clone expanded repeats with this method as well as perform SP-PCR with minor modifications to the standard protocols. Since the introduction of this technology in our laboratory, however, we have found that for some loci, we had to redesign primers. In addition to the transgene presented here, we have used the method to amplify the *DMPK* locus from myotonic dystrophy type 1 patient-derived lymphoblastoid cell lines. Thus, this approach to avoid carry-over contamination in PCRs is broadly applicable to expanded CAG repeats and possibly to other expanded microsatellite diseases. Moreover, we expect it to be useful for molecular diagnostic testing of expanded TNR repeat diseases.

## Materials and Methods

### Cell culture

GFP(CAG)_X_ cells used here are described elsewhere^16^,^22^. They were propagated in DMEM supplemented with 10% fetal bovine serum, 100 U ml^−1^ penicillin, 100 μg ml^−1^ streptomycin, 15 μg ml^−1^ blasticidine and 150 μg ml^−1^ hygromycin and were maintained at 37°C in a 5% CO_2_. They tested negative for mycoplasma at the beginning of our experiments using the MycoAlert Detection kit (Lonza LT07-218) and at the end using the service from GATC Biotech AG (Constance, Germany). The nickase treatment was done as before^16^, with GFP(CAG)1_01_ cells transfected with either pPN10, an empty plasmid, or pPN10-gCTG together with a vector expressing the Cas9 D10A nickase variant using Lipofectamine 2000 (Life Technologies) on day 1 and again on day 5 and 8. On day 12, we harvested the cells and found that those transfected with pPN10-gCTG used for the SP-PCR in Fig. 5 had an increase of 2.6 fold in the brightest 1% of the cells compared to pPN10-transfected cells. GFP intensities were determined using an Accuri C6 flow cytometer from BD.

### Bacterial strains

The *ung*^-^9 allele was made by amplifying the spectinomycin resistance gene from pENTR223.1 with oVIN-1448 and oVIN-1449 (see Table 4 for primer sequences) and transforming the resulting products into DH5α cells harbouring pKM208. This strain and plasmid were a gift of Kenan Murphy (Addgene plasmid # 13077). pKM208 contains the Red ***λ***phage proteins under a lactose-inducible promoter, a temperature sensitive origin of replication, and a gene expressing β–lactamase^23^. We followed a previously published protocol^20^, except that we generated electrocompetent cells using our standard laboratory protocol from cells grown to exponential phase at 30°C in LB supplemented with 100 ugml^1^ ampicillin and induced with 0.1 mM IPTG. After electroporation, the cells were selected at 37°C on LB agar plates containing 100 μg ml^−1^ of spectinomycin. The bacteria were then tested for the loss of ampicillin resistance (and thus loss of pKM208) at 30°C. Colony PCR was used to confirm the loss of *ung* and the gain of the spectinomycin resistance gene both 5’ and 3’ of the insertion site using primers oVIN-1451 and oVIN-1452 for the 5’ and oVIN-1450 and oVIN-1453 for the 3’ end. DH5α *ung^+^* control strains were made by curing the strain containing pKM208 obtained from Addgene by streaking the cells twice sequentially on LB agar plates incubated at 37°C and then selecting colonies that could not grow on LB agar plates containing 100 μg ml^−1^ ampicillin at 30°C. Competence was calculated by transforming a known amount of pUC19 and counting the resulting colonies on LB agar plates containing 100 μg ml^−1^ ampicillin.

**Table 4:**
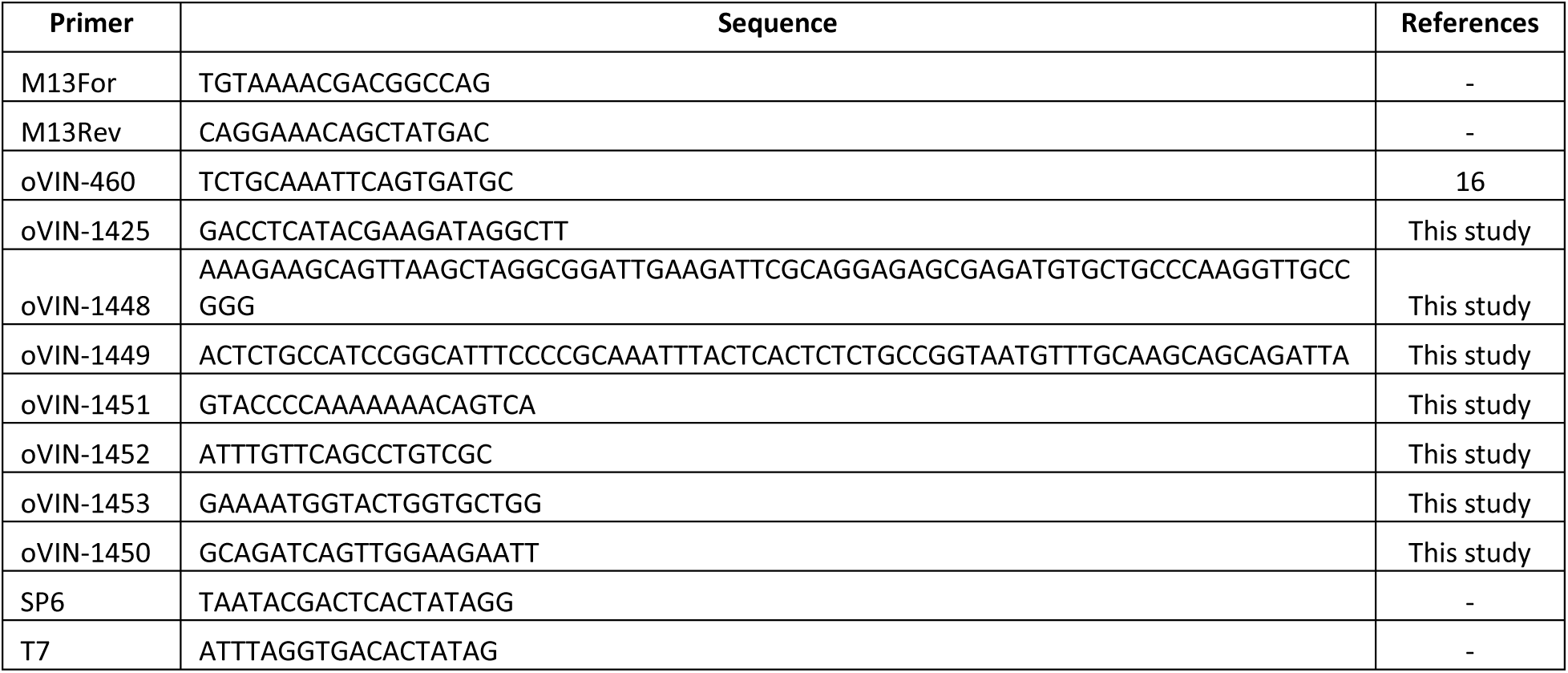
Primers used in this study

### Ung treatment, PCR amplification, and sequencing

The standard Ung was obtained from Thermo Fisher Scientific, whereas hlUng was from Roche. The polymerase used was the Phusion U Green Hot Start DNA Polymerase (Thermo scientific). We used 0.5 U of the Ung and 2 U of the Taq Polymerase in our reactions. The primers used to amplify the repeat tract from the GFP locus were oVIN-460 and oVIN-1425. The PCR program was as follow: 50°C for 2 min and 5 min at 95°C for the standard Ung, or 20°C for 10 min followed by 7 minutes at 95°C for hlUng. Then there were 35 cycles at 95°C for 30 s, 60°C for 30 s, and 72°C for 1 min 30 s. Ung was neutralized by the addition of Proteinase K (Promega) immediately after the PCR run to a final concentration of 100 ugml^−1^ and incubated at 37°C for 1 h. Sanger sequencing was done by Microsynth AG (Balgach, Switzerland).

### Molecular cloning

All plasmids used are in Table 5. For cloning the SUMO1 cDNA, we used pDONR-SUMO1 as PCR template using M13for and M13rev with hlUng and stopped the reaction as above. The amplicons were run on an agarose gel, the band was cut out and purified using the Nucleospin Gel and PCR clean up kit from Macherey-Nagel. The isolated DNA was quantified using a Nanodrop spectrophotometer. 220 ng of this insert DNA was used with the Zero Blunt Topo kit (Thermo Fisher Scientific). We diluted the ligation products 4-fold and used 2 μl of the reaction to electroporate 40 μl of bacteria, which were then plated onto LB agar plates containing 50 μg ml^−1^ of kanamycin. Colony PCRs were done with the SP6 and T7 primers, sequencing of the resulting plasmids was done with SP6.

**Table 5:** Plasmids used in this study.

| Plasmid | Content | References |
| --- | --- | --- |
| pKM208 | Ampicillin resistance, temperature sensitive origin of replication, Lambda RED proteins for recombineering. | 23 |
| pVIN-109 | GFP transgene containing 40 CAG repeats. Ampicillin resistance. | This study |
| pVIN-117 | GFP transgene containing between 40 and 89 CAG repeats and a modified intron to be described elsewhere. | This study |
| pPN10 | Kanamycin resistance, U6 promoter, guide RNA scaffold without a target sequence | 16 |
| pPN10-gCTG | Kanamycin resistance, single-guide RNA with 6 CTG repeats as target sequence | 16 |
| pDONR-SUMO1 | SUMO1 cDNA, Kanamycin resistance. | This study |

The degradation assays performed in Fig. 3 were done using 1 μg of plasmid DNA and 0.5U of Ung. They were incubated at 50°C for 30 min in PCR buffer. The amplicons used as controls for the degradation were the SUMO1 cDNA amplified as to make inserts for cloning. The PCRs contained dUTPs. We used the full content of a 50 μl PCR per reaction.

For cloning of the expanded repeats, we used the same protocol as for SUMO1 cDNA cloning, but the primers were oVIN-1425 and oVIN-460 and the template was pVIN-117, which harbours the GFP reporter assay with an unstable repeat tract containing up to 89 repeats. The resulting plasmids were sequenced with oVIN-460. The sequences were analyzed blindly of the genotype of the cells and of the nucleotide present in the insert. Of note, we found that sequencing in the other direction led to poorer results presumably due to the higher stability of CTG-containing secondary structures versus CAG-containing ones^24^.

### SP-PCR

The SP-PCR protocol was derived from^25^,using the primers and conditions used in^16^for dTTP-containing blots, with the exception that we used dUTP to generate the probe. A full step-by-step protocol is found in the supplementary material. *Statistics*

We used a two-tailed Student’s t-test that assumes equal variance to compare cloning efficiencies. When comparing the distribution of repeat size in Table 1 and Fig. 4, we used a two-tailed Wilcoxon U-test since the repeat size distributions were not normal. Nevertheless, the same conclusions were obtained with a two-tailed Student’s t-test. Statistical tests were performed using R version 3.1.3.

## Acknowledgements

We would like to thank Onya Opota for the initial suggestion of using Ung and dUTP in our PCRs. We also thank him, Fisun Hamaratoglu, and members of the Dion lab for critical reading of the manuscript. We are grateful to Yashashvi Singh Bhugowon for technical help and for blinding the sequencing data. This work was supported by a professorship from the Schweizerischer Nationalfonds zur Förderung der Wissenschaftlichen Forschung (#144789 to V.D.) and by Gebert Rüf Stiftung and UNISCIENTIA STIFTUNG within the programme «Rare Diseases – New Approaches» (GRS-060/14 to V.D.).

### Author contributions

L.A. and V.D. designed the experiments; L.A. performed them. V.D. performed the statistical tests and wrote the paper.

